# Reversible adhesion by type IV pili leads to formation of irreversible localized clusters

**DOI:** 10.1101/2022.01.24.477649

**Authors:** María Victoria Pepe, Celeste Dea, Camila Genskowsky, Darío Capasso, Adriana Valeria Jäger, Fernando Peruani, Arlinet Kierbel

## Abstract

Despite the fact a fundamental first step in the physiopathology of many disease-causing bacteria is the formation of long-lived, localized, multicellular clusters, the spatio-temporal dynamics of the cluster formation process, particularly on host tissues, remains poorly understood. Experiments on abiotic surfaces suggest that the colonization of a surface by swimming bacteria requires i) irreversible adhesion to the surface, ii) cell proliferation, and iii) a phenotypic transition from an initial planktonic state. Here, we investigate how *Pseudomonas aeruginosa* (PA) infects a polarized MDCK epithelium and show that contrary to what has been reported on the colonization of abiotic surfaces, PA forms irreversible bacterial clusters on apoptotic epithelial cell without requiring irreversible adhesion, cell proliferation, or a phenotypic transition. By combining experiments and a mathematical model, we reveal that the cluster formation process is regulated by type IV pili (T4P). Furthermore, we unveil how T4P quantitatively operate during adhesion on the biotic surface, finding that it is a stochastic process that involves an activation time, requires the retraction of pili, and results in reversible adhesion with a characteristic attachment time. Using a simple kinetic model, we explain how such reversible adhesion process leads to the formation of irreversible bacterial clusters and quantify the cluster growth dynamics.

---

The early stages of many infection processes, which remain poorly understood, require bacteria to localize suitable host tissues where to anchor and form bacterial multicellular structures such as biofilms^1^. Often, the tissue colonization starts with the formation of localized bacterial clusters^2–6^. Once within mature multicellular structures and biofilms, bacteria are embedded in the extracellular matrix, which can be self-produced and/or formed with material acquired from the host tissue^7^, and exhibit resistance to flows and importantly, an increased tolerance to antibiotics and immune system responses.

For technical reasons, as well as for its relevance in industrial applications, bacterial colonization and biofilm formation have been investigated on abiotic (and generally spatially homogeneous) surfaces^8–14^. 2

It has been observed that an initial population of planktonic bacteria undergoes various phases before actual colonization of the surface^9,10,12^. In the initial phase that elapses for several hours, the overwhelming majority of bacteria remains swimming in the fluid, and attach only reversibly to the surface^8–10,12^. During this phase, it is believed that the bacterial population, over repeated cycles of surface sensing and detachment, becomes progressively adapted for irreversible surface attachment^11,12^. This is evidenced in the next phase of the process by a sudden exponential growth of the surface bacterial population that leads to a quick surface coverage that involves irreversible attachment, bacterial proliferation, and extracellular matrix production^8,12^. For *Pseudomonas aeruginosa* (PA) on abiotic surfaces^12,15,16^, the initial reversible-attachment phase elapses for 20 hours. It is only after this initial period that irreversible attachment leads to the formation of nascent bacterial clusters^15^.

The colonization of biotic surfaces, on the other hand, remains largely unexplored. Infection experiments with polarized MDCK cells have revealed that PA is able to form bacterial clusters primarily on apoptotic cells shedding from the epithelium. These clusters reach their final size in minutes and remain stable for hours^17,18^. How PA form such irreversible clusters has not yet been known. Here, we investigate the growth dynamics of these PA clusters and the statistics of the bacterial adhesion times. By combining experiments and a mathematical model, we find that the cluster formation process is regulated by type IV pili (T4P), which are hair-like appendages that can be rapidly extended and retracted to generate active forces to move or adhere^1920^. Furthermore, we reveal how T4P quantitatively operate during adhesion on the apoptotic cells, finding that it is a stochastic process that involves an activation time, requires the retraction of pili, and results in reversible adhesion. In addition, we quantify the cluster growth dynamics and explain how such reversible adhesion process leads to the formation of irreversible bacterial clusters that are arguably the precursors of a full-scale tissue infection. In short, our study shows that irreversible bacterial cluster formation in PA on biotic surfaces does not require irreversible adhesion, cell proliferation, or any phenotypic transition, in sharp contrast to what has been reported for PA on abiotic surfaces.

## Results

### Features of formed bacterial clusters

On the polarized MDCK epithelium, PA forms bacterial clusters, on sites of apical extrusion of apoptotic cells, which we refer to as clusters or aggregates. In the span of minutes, free-swimming bacteria are recruited on the surface of those apoptotic cells^17,18^. Round-shaped bacterial aggregates of approximately 10 microns diameter are observed after infecting MDCK monolayers with PA strain K for one hour (Figs. 1A and B). We investigate how PA attach to apoptotic cells, by measuring the angle between the longitudinal axis of the bacterium and the tangent of the cell surface; see Fig. 1C, where only bacteria on the focal plane are taken into consideration. We find that bacteria attach to the host cell with the cell body parallel to the normal vector of the cell membrane. Note that this spatial arrangement allows bacteria to densely cover the host cell. In previous studies, interaction with a surface via the cell pole has been associated with a reversible attachment, while irreversible attachment has been thought to require the cell to orient parallel to the surface^21^. We recall that WT PA harbors one flagellum and a reduced number of T4P, located at the bacterium poles. To visualize the flagellated pole in live bacteria the monolayers are infected with PA expressing chemotaxis protein CheA bound to GFP (CheA-GFP). CheA has a unipolar localization pattern at the flagellated pole^22–24^. We find 74% of the bacteria attached to apoptotic cells by the pole opposite to CheA (Fig. 1D) and therefore opposite to the flagellum, indicating that there is a preferential orientation and discouraging the idea that the flagellum plays a role of an adhesin in this system. Nonetheless, flagella are essential for aggregate formation. The aflagellated mutant ΔfliC (the gen flic encodes the major component of the flagellum) is unable to form aggregates (Supplementary Figure 1). This is expected as bacteria reach apoptotic cells by swimming.

**Fig. 1.**
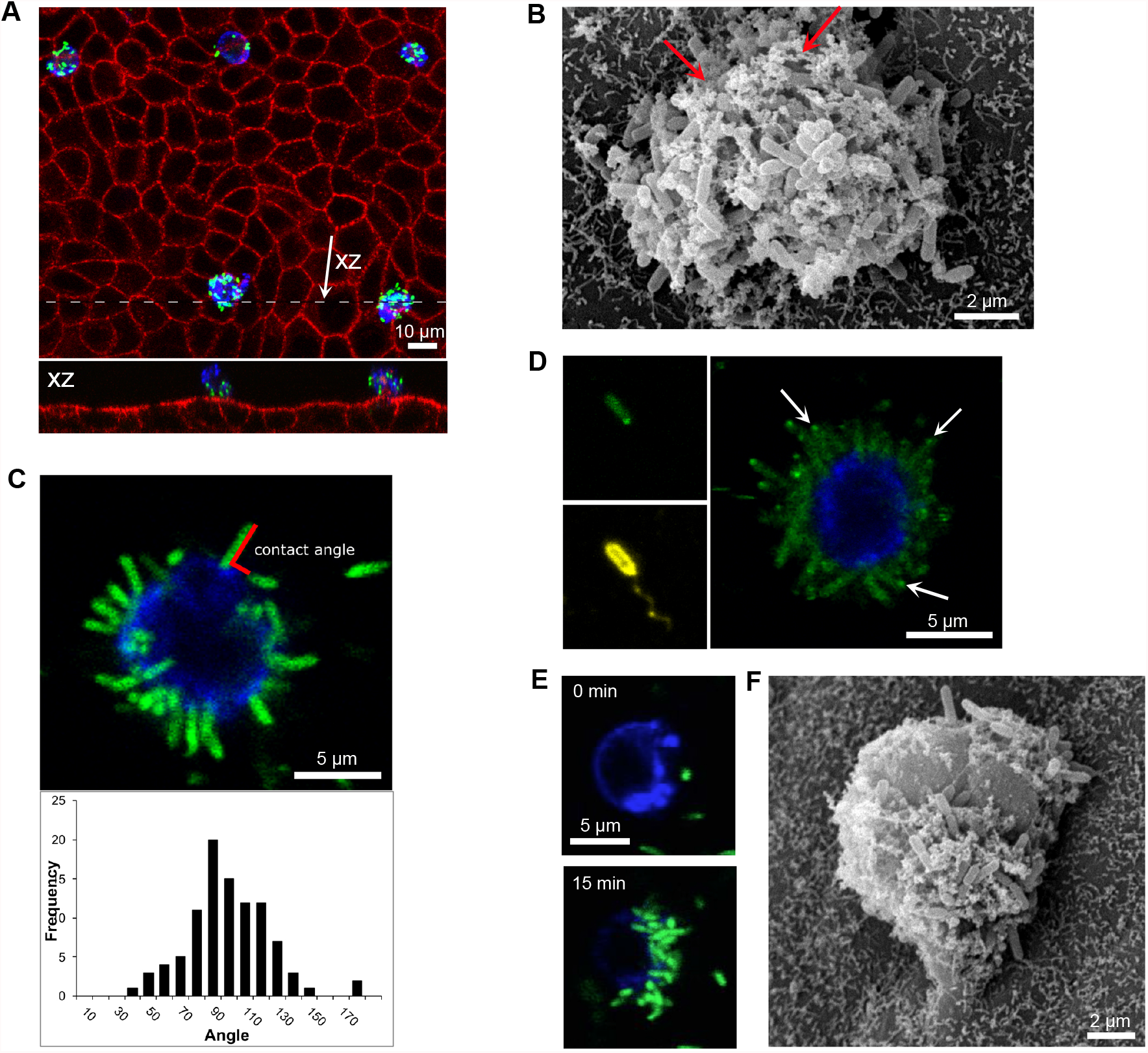
Morphology of formed PA aggregates. (A, B and F) Transwell grown MDCK monolayers were infected with PA, incubated for 1 h and fixed. (A) Top view and orthogonal section (upper and lower panel respectively) showing a confocal micrograph of a monolayer with several extruded apoptotic cells with adhered bacteria. After infection with PA-GFP (green), samples were labeled with Annexin V-Alexa 647 (blue), fixed, permeabilized and stained with phalloidin-rhodamine for F-actin (red). (B and F) Scanning electron micrographs. (B) Bacterial aggregate. Arrows indicate apoptotic host cell material. (C-E) Time-lapse confocal microscopy images. Annexin V: blue. (C) The monolayer was infected with PA-GFP (green). The angle between the longitudinal axis of bacteria and the tangent of the cell surface was measured as indicated (upper panel). The 93% of the angles fell between 45 and 135 degrees (lower panel) showing that bacteria attach by the pole. (D) CheA was used as a reporter of the flagellar pole (left panels show a fixed bacterium expressing CheA-GFP (green) and stained with an anti-PA antibody that labels the flagellum (yellow)). Right panel: time-lapse image showing bacteria attach to apoptotic cells by the pole opposite the flagellum. (E) Micrographs show the same apoptotic cell at the beginning (upper panel) and 15 minutes after infection (lower panel) with PA-GFP (green). Bacteria adhered to zones of more intense annexin V labeling. (F) Bacteria attach to zones of the surface with vesiculated morphology.

Biotic surfaces can display complex and heterogeneous topographies. Particularly, the plasmatic membrane of apoptotic cells suffers dramatic changes as the apoptotic process evolves. Upon infection, most extruded apoptotic cells are fully covered with bacteria. However, in some apoptotic cells it is observed that bacteria are distributed heterogeneously over the membrane, with patches that are covered with bacteria and other areas that are bacteria-free. We investigate whether there are detectable differences between membrane areas occupied and unoccupied by bacteria. Notably, in areas where bacteria attach, AnnexinV labeling is more intense (Fig. 1E and Supplementary Video 1). The quantification indicates that there exists a positive correlation between fluorescence intensity and bacterial number (Spearman Correlation’s coefficient r = 0.77, p <0.05). Staining with a general membrane marker displays a similar result (Supplementary Figure 2), suggesting that bacterial attachment occurs in zones of increased membrane surface availability. Then, infected and uninfected samples are analyzed by Scanning Electron Microscopy. Fig. 1F shows an extruded apoptotic cell with heterogeneous areas of adhered bacteria. Notably, the surfaces covered with bacteria are filled with membrane-enclosed microvesicles or small apoptotic bodies. In contrast, the bacteria-free membrane is smooth. Importantly, extruded cells of vesiculated morphology were also present in uninfected samples. And PA adheres all over the surface of apoptotic cells that are fully covered by microvesicles. This kind of cell surface has an irregular topography (Supplementary Figure 3). In recent years, surface roughness and topography have been found to be critical to bacterial adhesion^13,14^.Taken together, our results indicate that PA attaches to extruded apoptotic cells vertically, by the pole opposite the flagellum, and demonstrate preference for cell surfaces with an irregular topography.

**Fig. 2.**
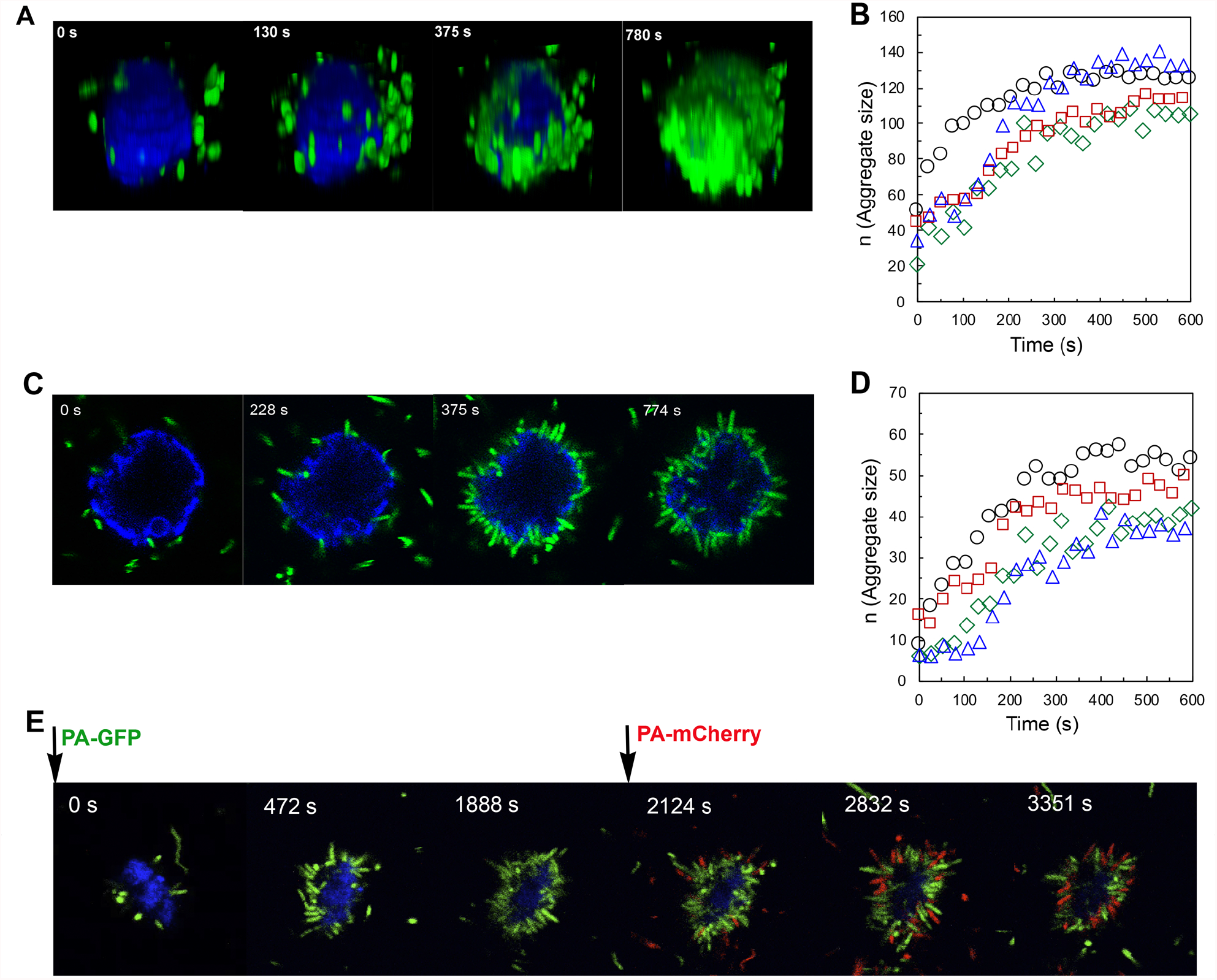
Formation of aggregates on apoptotic cells extruded from a monolayer. (A,C and E) Time-lapse confocal imaging of PA-GFP (green) adhering on apoptotic cells (blue). (A) 3D reconstructions of successive z-stacks (C) Snapshots of the equatorial plane of the cell. (B and D) Growth curves of four different experimental aggregates. (B) The number of bacteria (aggregate size, denoted by *n*) was obtained from the entire z-stack. (D) The number of bacteria found on the equatorial plane. (E) Snapshots of the equatorial plane in an experiment where initially the monolayer was inoculated with PA-GFP (green) and after 30 min PA-mCherry (red) was added

### Temporal dynamics of aggregate formation

Immediately after wild-type (WT) PA is released, apoptotic cells start to be visited by bacteria and aggregate formation begins. We quantify the growth of the cluster by counting the number of bacteria in three dimensions as well as at the equatorial plane of the apoptotic cell as shown in Fig. 2 A and C and Supplementary Video 2. While both methods provide comparable information on the cluster dynamics, see Fig. 2B and D, the latter allows a faster acquisition rate. Once clusters are formed, they remain stable in size for at least 3 hours. The observed dynamics leads to the formation of irreversible bacterial clusters, in the sense that once formed, the cluster size remains roughly constant over time, and thus the cluster is long-lived. However, careful inspection of the data shows that during cluster growth bacteria forming the cluster often detach and swim away from it, leaving an area of the apoptotic cell membrane vacant. This vacant membrane area is exposed to free-swimming bacteria, and thus at some later time becomes occupied again. The bacterial attachment-detachment process continues even in fully formed clusters. The reversible character of the adhesion process can be experimentally evidenced using two differentially labeled populations of PA, as shown in Fig. 2E and Supplementary Video 3. Note that the reversible adhesion implies that clusters are dynamic structures. How can we characterize the observed growth dynamics and understand the emergence of irreversible, dynamical structures, when bacteria reversibly attach and detach from it? In order to quantify the cluster formation process, we focus on the dynamics of a small membrane area of the apoptotic cell that can be either vacant or occupied at most by a bacterium. Our first task is to characterize from the experiments the times during which the small membrane area is occupied; for details on the computation of these times see *Materials and Methods*. The distribution of these times is presented in Fig. 3A in the form of a survival curve *S*(*t*), which indicates the probability of observing a dwelling time greater than or equal to *t*. Note that *S*(*t*) for WT-PA in Fig. 3A is not given by a simple exponential. Thus, if we attempt to mathematically model the dynamics assuming two states for the small membrane area – e.g. state 0 for vacant and state 1 for occupied, and transition rates *r*_01_ and *r*_10_ for transitions 0 → 1 and 1 → 0, respectively – we will fail to explain the experimentally obtained distribution *S*(*t*). For a two-state Markov chain as described above, the survival curves associated to staying in state 0 and 1 are both, single exponential. And thus, in order to explain the measured distribution of times for WT PA, we are mathematically forced to assume – in order to consider a larger family of functional forms, including the experimental ones – the existence of (at least) three states: 0, 1, and 2. Furthermore, states 1 and 2 necessarily correspond, both of them, to occupied states of the membrane area. But, what is the interpretation of these mathematically postulated states? The existence of these two occupied states suggest two different types of (transient) membrane adhesion. In order to shed light on the role of T4P, we analyze experiments with T4P mutants: i) non-piliated ΔPilA mutant – PilA is the major pilin subunit – and ii) hyperpiliated ΔPilT mutant – PilT is the molecular motor that mediates pilus retraction. Thus, ΔPilT mutants are unable to retract their pili. From the comparison between these mutants, we learn that a) clusters only emerge in WT (see also Supplementary Figure 4 and Supplementary videos 4 and 5), b) two occupied states are required to account for dwelling-time distributions of WT and ΔPilT, i.e. for bacteria displaying T4P, and c) the dwelling-time distribution is a single exponential for non-piliated ΔPilA only. In consequence, state 2 is only present for bacteria equipped with T4P, i.e. WT and ΔPilT, and thus it can be associated to T4P-mediated adhesion. The dynamics among these states – see Fig. 3B – is given by:

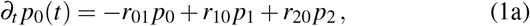

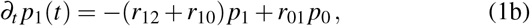

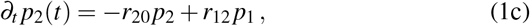

where *p*_*i*_(*t*) is the probability of finding the small membrane area in state *i* and *r*_*i j*_ are the transition rates between state *i* and *j*. From Eq.(1), we compute *S*(*t*) as a first-passage time problem that indicates for how long the system remains between state 1 and 2 before transitioning to 0, which reads:

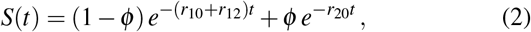

where *ϕ* = *r*_12_*/*(*r*_12_ + *r*_10_ − *r*_20_). By applying Eq. (2) to describe the distributions in Fig. 3A, we find *r*_10_ = 0.28 ± 0.01 s^−1^, *r*_12_ = 0.03 ± 0.01 s^−1^, and *r*_20_ = 0.004 ± 0.0009 s^−1^ for WT, and *r*_10_ = 0.25 ± 0.01 s^−1^, *r*_12_ = 0.02 ± 0.01 s^−1^, and *r*_20_ = 0.055 ± 0.002 s^−1^ for ΔPilT; further details in *Material and Methods*. On the other hand, for ΔPilA data *r*_12_ = 0 and thus the model becomes effectively a twostate Markov chain, with only state 0 and 1 participating into the dynamics, and *S*(*t*) reduces to 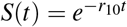, obtaining *r*_10_ = 0.23 0.02 s^−1^. Note that the main different between the rates of WT and ΔPilT is observed in *r*_20_ that is 10 times larger for ΔPilT, which implies that dwelling times are expected to be in average one order of magnitude longer in WT. The similarity of the obtained values *r*_10_ in experiments with WT, ΔPilT mutants, and ΔPilA mutants suggests that the transition 0 → 1 involves the same mechanism for WT and these mutants, which is evidently unrelated to T4P. In summary, the transition from 1 → 2 observed in WT and ΔPilT mutants indicates that adhesion mediated by T4P is a stochastic process that requires not only the presence of pili, but also the capability of retraction it to achieve long adhesion times. It is worth stressing that Eq. (1) is the simplest 3-state Markov chain consistent with the experimental data: transition rates *r*_02_ and *r*_21_ can be also included in the description in order to allow all possible transitions, however, these extra two parameters do not improve the goodness of the fit; and thus including them leads to over-fitting. For further details on the derivation of Eq. (2) and fitting procedure, see *Materials and Methods*. Now, we focus on the growth of the cluster. We consider the probability *P*(*n, t*) of finding *n* bacteria on the apoptotic cell at time *t*, assuming the cell contains *N* statistically independent small membrane areas. Under these assumptions, we approximate the evolution of *P*(*n, t*) by the following master equation:

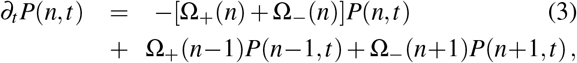

where Ω_+_(*n*) = *α*_+_(*N* − *n*) and Ω_−_ (*n*) = *α*_−_ *n*. The rate *α*_+_ is directly *α*_+_ = *r*_01_ and describes how frequently swimming bacteria arrive at a vacant membrane area. And thus, this rate depends on bacterial motility as well as on bacterial density; the simplest assumption is that *α*_+_ ∝ *C*, with *C* the inoculated bacterial concentration. On the other hand, *α*_−_ depends on intrinsic properties of the bacterium, i.e. on its adhesion capacity to the apoptotic cell membrane, and is given by inverse of the average time a bacterium remains on the cell membrane, related to *S*(*t*) by:

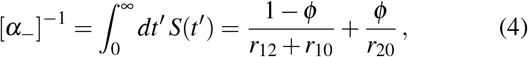

implying, 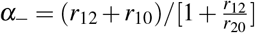. The solution of Eq. (3) with the provided definitions of Ω_+_(*n*) and Ω_−_(*n*) and using as initial condition that at *t* =0 there is no bacteria on the cell – i.e. *P*(*n* = 0, *t* = 0) = 1 and *P*(*n, t* = 0) = 0 for *n >* 0 – reads:

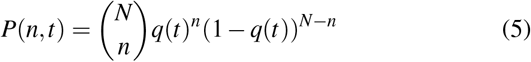

with 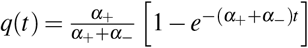; see Fig. 3C and D. The binomial nature of Eq. (5) implies that ⟨*n*⟩ (*t*)=∑_*n*_ *n P*(*n, t*) = *q*(*t*)*N*. The advantage of the approximation given by Eq. (3) is that it allows us to show that the growth of the cluster can be conceived as a biased random walk in the cluster-size space: the walker can move from position *n* to either *n* −1 (after the detachment of a bacterium) or *n* + 1 (if a bacterium attaches to the cell). The ratio of the transition probabilities *n* → *n*+1 and *n* → *n* − 1 provides an idea of the local bias of the walker, which depends on *n* as well as on the ratio *α*_+_*/α*_−_; Fig. 3D. At small values of *n*, the large availability of vacant sites, i.e. *N* − *n*, favors a bias toward large *n*-values, and the opposite happens for large values of *n*. If rates *α*_+_ and *α*_−_ are identical, then the walkers moves to, and remains around, *n*_∗_ = *N/*2, but in general *α*_+_*/α*_−_ ≠1, and the equilibrium position corresponds to *n*_∗_ = *N/*(1 + *α*_−_*/α*_+_). We note the critical dependency of *α*_−_ with *r*_12_. In the limit of large *r*_12_ values, *α*_−_ ∼*r*_20_, while for *r*_12_ → 0, *α*_−_ → *r*_10_. Since *r*_20_ ≪ *r*_10_, the equilibrium position for WT, equipped with a fully functioning T4P, is expected to be much larger than the one for ΔPilA and ΔPilT mutants. The analogy with the biased random walk allows us to conceptually understand the emergence of an irreversible dynamics for cluster growth out of the reversible, attachment-detachment action of individual bacteria. However, the approximated temporal evolution of *P*(*n, t*) given by Eq. (3) assumes that transitions from *n* → *n*+1, *n* → *n* − 1, etc are characterized by exponentially distributed times, which is certainly not true as evidenced by the distribution of dwelling times, Fig. 3A. Nevertheless, it is possible to obtain an exact solution of the original problem using that at every time *t* the probability is given by the binomial distribution 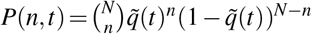, with 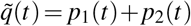, where *p*_1_(*t*) and *p*_2_(*t*) are the solutions of Eq. (1) with initial condition *p*_0_(*t* = 0) = 1 and *p*_1_(*t* = 0) = *p*_2_(*t* = 0) = 0; see *Material and Methods* for explicit expressions. In Fig. 3E, the exact and approximate solution are used to describe temporal evolution of cluster size on an apoptotic cell. The value of *r*_01_ is adjusted via the nonlinear least squares method obtaining *r*_01_ = 0.037 ± 0.002s^−1^; for further details see *Material and Methods*. Note that the approximate solution fails to describe the temporal evolution towards the equilibrium cluster size, which indicates that considering three states is key to achieve a faithful quantification of the temporal dynamics; Fig. 3E.

**Fig. 3.**
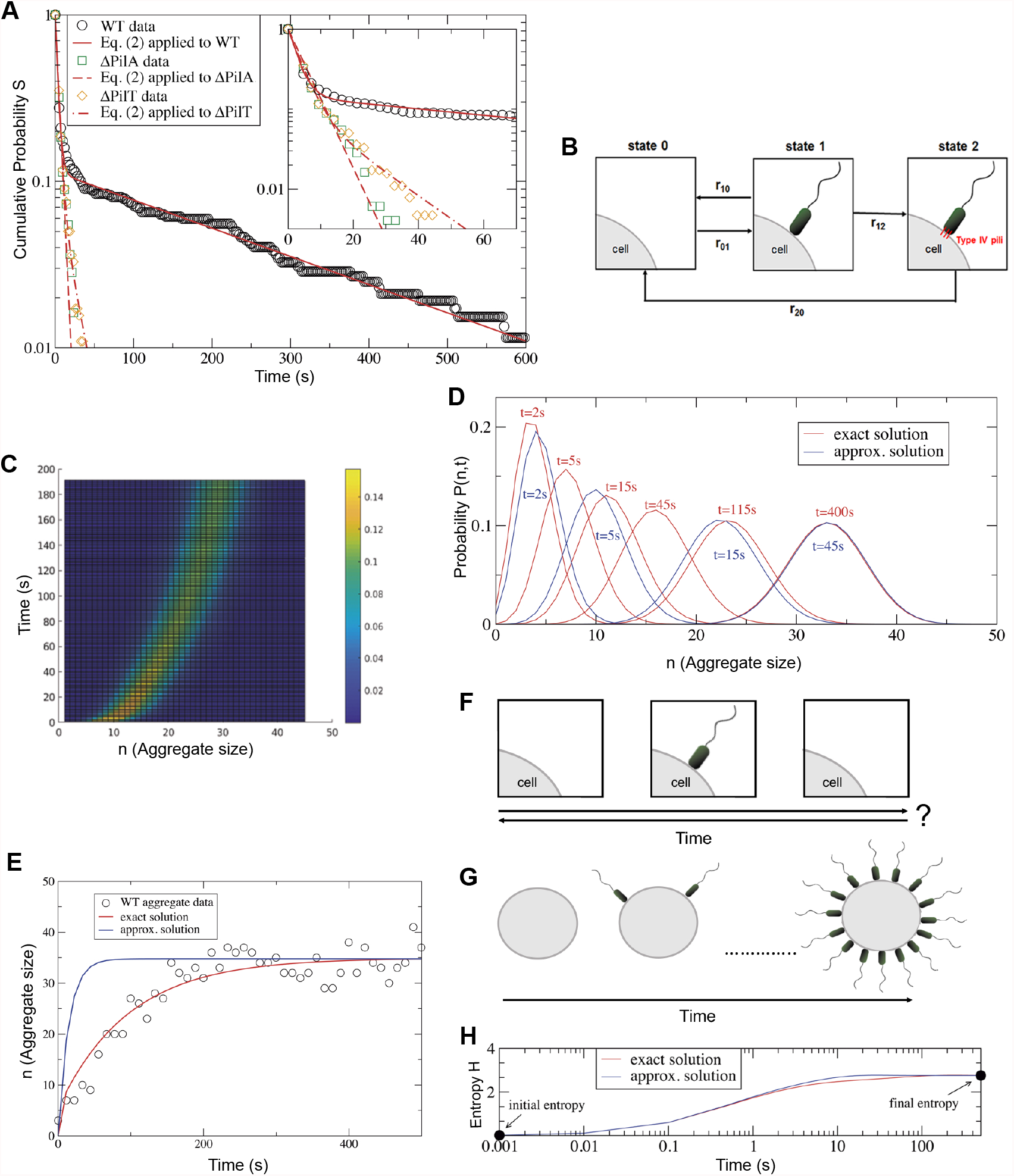
Growth dynamics of the aggregate. (A) Semi-log plot of the cumulative distribution of bacterial dwelling times on the cell membrane. Circles correspond to WT data, squares to ΔPilA data, and diamonds to ΔPilT data, while the solid, dashed, and dot-dashed curves to Eq. (2) applied to WT, ΔPilA, and ΔPilT data, respectively. The inset displays the distributions for short dwelling times, in the range [0, 70]. (B) Scheme of the three-states model, see Eq. (1). For *r*_12_ = 0, the dynamics reduces to a 2-state model, with only states 0 and 1. (C) Temporal evolution of the probability *P*(*n, t*) (color coded) of finding that at time *t* the aggregate size is *n*; see Eq. (5). (D) Comparison of the exact (red) and approximate (blue) solution of *P*(*n, t*), evaluated at various times *t*. (E) Aggregate size *n* vs time. Circles correspond to the growth of an experimental aggregate, while the red and blue curve correspond to exact and approximate solution of *P*(*n, t*), respectively. Schemes (F) and (G) illustrates that the vacant-occupied dynamics of a small cell membrane area does not convey information about the arrow of time, (F), while from the temporal evolution of the aggregate is possible to identify it (G). (H) The increase in entropy *H*, Eq. (6), puts in evidence the arrow of time and the irreversible character of the growth of the aggregate.

## Discussion

The dynamics of a small cell membrane area is such that it is at times vacant and at times occupied by a bacterium, undergoing a state cycle between vacant and occupied. This implies that if the dynamics of this small membrane area is recorded in a video, and is shown to us, we will not be able to determine whether it is played forwards or backwards, i.e. we will not be able to identify the arrow of time; see Fig. 3F. On the other hand, if we watch a video of the evolution of the whole cluster, we can easily determine whether the video is played forward or backward, and thus the arrow of time (and irreversibility) becomes apparent; Fig 3G. The analogy with the biased random walk in cluster size space has allowed us to mathematically understand the emergence of irreversibility out of a reversible dynamics at the level of individual bacteria. A formal way to put in evidence the irreversible character of cluster growth is to define the (Shannon) entropy of this structure as:

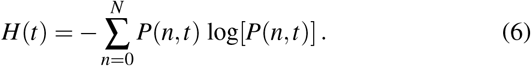

The temporal evolution of this quantity, which scales with the apoptotic cell size, is displayed in Fig. 3H that shows that *H* starts at a low level and reaches a final larger entropy value as is expected in an irreversible process. As the system will not spontaneously (in average) decrease its entropy at a later time, the cluster will not disintegrate. Note that the behavior of *H*(*t*) is almost identical for the exact and approximated solution of *P*(*n, t*), implying that the cluster dynamic is irreversible for both. Mathematically, the approximated solution given by Eq. (3) is based on an effective reduction of the dynamics to two states, while the exact solution is based on three states. This indicates that mathematically the use of three states – which at microscopic level, according to Eq.(1), involves entropy production – is not a necessary condition to obtain an irreversible cluster dynamics. Considering three states and their interplay is, however, essential, not for irreversibility, but to obtain an accurate description of dwelling times and of cluster growth, and unveils fundamental information on T4P-adhesion dynamics. In particular, state 2 is required for an accurate description of adhesion times of bacteria equipped with T4P. The presence of this state is a necessary, but not sufficient condition to observe long adhesion times and cluster formation (cf. WT, ΔPilT, and ΔPilA). In addition to the presence of T4P, the capacity of retracting it is necessary, which suggests that anchoring on the membrane occurs during retraction, in a dynamics reminiscent of catch-bond adhesins^25^. Furthermore, the three-state model allows to infer how T4P mediated adhesion works on the cell membrane: first the bacterium needs to reach the cell membrane (transition 0 → 1), once in contact with the cell membrane, the T4P-adhesion can be triggered in an average of 33s (transition 1 → 2), and remains activated an average time of 4.3min (transition 2 → 0). These findings are inline with recent results obtained by Koch et al.^26^ that indicate that the T4P-apparatus operates by stochastically extending and retracting pili, and observe that these events are not triggered by surface contact. On the other hand, we recall that PA is temporally attached upright by the pole opposite to the flagellum; Fig. 1D. It is worth mentioning that Schniederberend et al.^27^ found that when the attachment is, contrary to what is reported here, mediated by the flagellum, irreversible adhesion is induced and PA ends up laying horizontally on the surface. This suggests that adhesion by the pole opposite to the flagellum is characteristic of transient adhesion mediated by T4P. We note that the stochastic character of the adhesion process has also been evidenced in other bacterial systems, where interestingly active adhesion has been described by a two-step process^28,29^, an observation that suggests possible universal adhesion behaviors.

Finally, we speculate on the functionality of the observed dynamical multicellular structures. An often invoked advantage of bacterial clustering is that it allows bacteria to cooperate by sharing “public goods”. This can occur by collectively secreting enzymes into the surroundings in order to digest too large or insoluble materials^30,31^. As the amount of hydrolyzing enzymes increases, the local concentration of oligomers to uptake also does it^32^. Thus, it can be speculated that in clusters with a constant turnover of bacteria, the concentration of “public goods” (enzymes or signals) increases by allowing, overtime, a larger number of donors to participate.

## Methods

### Time-lapse confocal microscopy

*P. aeruginosa* K (PAK) strains WT and ΔFliC, ΔPilA and ΔPilT mutants (kindly provided by J. Engel) were used. For CheA localization experiments the plasmid pJN(cheA-gfp) was used^24^. For time-lapse microscopy studies MDCK cells were grown on 35 mm Glass bottom dishes (10^4^ cells per cm^−2^ were seeded and grown for 72 h to ensure polarization). Monolayers were washed with binding buffer, incubated with Alexa conjugated-Annexin V for 15 min (Annexin V binds to phosphatidylserine, which is located on the outer leaflet of the apopototic cell membrane) and then washed with MEM. Monolayers were then incubated in MEM supplemented with HEPES 20 mM. Microwell dishes were placed on the microscope stage and the stack dimensions were set up from top to bottom throughout one or a few apoptotic cells (10 confocal optical sections at 1 *μ*m intervals). Fluorescent bacteria were inoculated and immediately after image acquisition was started. This process was conducted at 25 C. Alternatively, only the equatorial plane of the cell was scanned. Images were recorded with a confocal laser-scanning microscope Olympus FV1000, using a PlanApo N (60X 1.42 NA) oil objective. The image size was 512 × 512 pixels. To measure “residence times”, monolayers were inoculated with a mix of *P. aeruginosa*-GFP and *P-aeruginosa*-mCherry (1:3, final MOI = 20). In these experiments images were acquired at 2.33 sec/frame. To track green bacteria and establish the attachment and detachment times, we used the MTrackJ plugin from the ImageJ software (National Institutes of Health, NIH, USA). MTrackJ plugin facilitates manual tracking of moving objects in image sequences. For image acquisition of fixed samples the image size was 1024 × 1024, and the z-stack interval 0.3 *μ*m. More details are provided in Supplementary Information.

### Mathematical model and fitting procedure

We provide details on the exact solution of Eq. (1), the derivation of *S*(*t*), and the fitting procedure.

*Exact solution of Eq. (1)*.*–* Let us first recast Eq. (1) as:

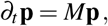

where **p** = (*p*_0_, *p*_1_, *p*_2_)^*T*^ and

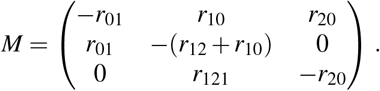

We use as initial condition: **p**(*t* = 0) = (1, 0, 0)^*T*^. The exact solution takes the form **p** = *c*_0_ exp(*λ*_0_*t*)**V**_0_ + *c*_1_ exp(*λ*_1_*t*)**V**_1_ + *c*_2_ exp(*λ*_2_*t*)**V**_2_. The eigenvalues are *λ*_0_ = 0, *λ*_1_ = (−*ω* − *β*) */*2, *λ*_2_ = (−*ω* + *β*) */*2, with 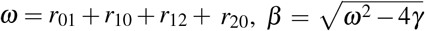, where *γ* = *r*_01_(*r*_12_ + *r*_20_) + *r*_20_(*r*_10_ + *r*_12_), their corresponding eigenvectors are:

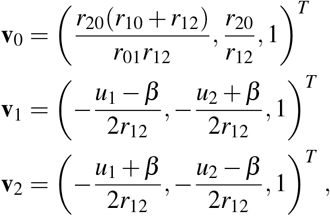

where *u*_1_ = − *r*_01_ − *r*_10_ + *r*_12_ + *r*_20_ and *u*_2_ = *r*_01_ + *r*_10_ + *r*_12_ − *r*_20_. Finally, the coefficients are 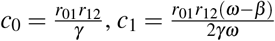, and 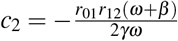. We stress that 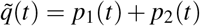, used to construct the exact solution *P*(*n, t*) in the main text, corresponds to this solution, and should not be confused with *S*(*t*).

#### Derivation of S(t)

From Eq. (1), *S*(*t*) is computed as a first-passage problem: assuming that at *t* = 0 the state is 1, we estimate for how long state remains between 1 and 2, before transitioning back to 0. The system to solve is:

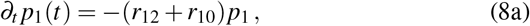

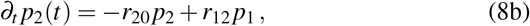

with initial condition *p*_1_(*t* = 0) = 1 and *p*_2_(*t* = 0) = 0. The survival probability *S*(*t*) is directly *S*(*t*) = *p*_1_(*t*) + *p*_2_(*t*), whose explicit solution is given by Eq. (2). It is important to stress that *S*(*t*) is not *p*_1_(*t*) + *p*_2_(*t*) of Eq. (1), but of Eq. (8) with the specified initial conditions.

#### Fitting procedure

For the analysis of the dwelling times, we have to consider that we are limited by the duration of the experiment. In consequence, there are dwelling events, where we observe the beginning, i.e. when the bacterium attaches to the membrane, but not the end of the event, i.e. when the bacterium detaches, since we arrive at the end of the experiment. We classify dwelling times in two categories: those where we have observed the beginning and the end of the event (uncensored data), and those where we have observed the beginning, but not the end, which we analyzed using the Kaplan-Meier method. The fitting of data is obtained by applying nonlinear least squares to the obtained analytical expressions. We find using Eq. (2) for uncensored data, *r*_10_ = 0.28s^−1^, *r*_12_ = 0.03s^−1^, and *r*_20_ = 0.004s^−1^ [*χ*^2^ = 0.007, *R*^2^ = 0.997], while for the Kaplan-Meier method, *r*_10_ = 0.29s^−1^, *r*_12_ = 0.09s^−1^, and *r*_20_ = 0.002s^−1^ [*χ*^2^ = 0.01, *R*^2^ = 0.98]; (Supplementary Figure 7). In ΔPilA data, *r*_12_ = 0 and *r*_10_ = 0.23s^−1^ [*χ*^2^ = 0.004, *R*^2^ = 0.998], and in ΔPilT data, *r*_10_ = 0.25 s^−1^, *r*_12_ = 0.02 s^−1^, and *r*_20_ = 0.055 s^−1^ [*χ*^2^ = 0.001, *R*^2^ = 0.999]. censoring data for ΔPilA and ΔPilT mutants is not necessary given the short duration of dwelling times. Finally, the description of aggregate growth is performed via *P*(*n, t*). All parameters, but *r*_01_ are determined by the dwelling time distribution. Using the set of rates corresponding to the uncensored data, we find *r*_01_ = 0.04s^−1^ [*χ*^2^ = 3075.7, *R*^2^ = 0.90], and with the ones for all data, *r*_01_ = 0.035s^−1^ [*χ*^2^ = 3535.3, *R*^2^ = 0.88]; (Supplementary Figure 5).

## Supplementary Information

### SI Materials & Methods

#### Antibodies and reagents

Anti-*P. aeruginosa* antibody (ab68538) was obtained from AbCam. Alexa-conjugated Annexin V, Phalloidin-Rhodamine and CellMask Deep Red were obtained from Thermo Fisher Scientific.

#### Cell culture and bacterial infection

MDCK cells (clone II, generously gifted by Dr. Keith Mostov) were cultured in MEM containing 5% fetal bovine serum. For time-lapse experiments, cells were grown on glass-bottom dishes with a 35 mm micro-well (MatTek Corporation). Around 10^4^ cells per cm-^2^ were seeded and then kept for 72 h in culture to ensure the formation of fully polarized monolayers. For studies with fixed samples, cells were grown on 12-mm transwells (Corning Fisher, 4.5×10^5^ cells per transwell) and used for experiments after 48 h in culture. Annexin V-Alexa-647 staining was done in binding buffer (10 mM HEPES, 140 mM NaCl and 2.5 mM CaCl2, pH 7.4).

Bacteria were routinely grown shaking overnight in Luria-Bertani broth at 37°C. Plasmids used were: pMP7605 (FEMS Microbiol Lett. 2010; 305(1):81–90), pBBR1MCS-5 + gfpmut3, pSV35 + pilA and pJN(cheA-gfp) (*Mol. Micro*, 90, 923– 938, 2013). Stationary-phase bacteria were co-incubated with epithelial cells at a MOI of 20 for confocal studies, and at a MOI of 60, for scanning electron microscopy studies.

#### Microscopy studies

To visualize CheA localization, bacteria carrying the CheA-GFP plasmid were allowed to adhere to polylysine-treated slides for 30 minutes at room temperature. Samples were fixed with 4% paraformaldehyde in PBS for 15 minutes, blocked with BSA 1%, and incubated overnight at 4°C with the Anti-P. aeruginosa antibody. To measure the number of bacteria per aggregate, transwell-grown MDCK-monolayers were infected with the indicated strains for 1 h (MOI: 20). Samples were labeled with Alexa conjugated-Annexin V, fixed, blocked, permeabilized with saponin 0.1%, stained with phalloidin for 60 minutes, and analyzed by confocal microscopy. For scanning electron microscopy, transwell-grown MDCK monolayers were infected with *P. aeruginosa* for 1 h. Samples were washed with 0.15M Sorensen buffer (0.056 M NaH2PO4, 0.144 M Na2HPO4 pH = 7.2) and fixed with 2.5% glutaraldehyde in 0.1M Sorensen buffer for 1 h at room temperature. Samples were then washed, and progressive dehydration was carried out. After critical point drying and gold sputtering, samples were analyzed with a Carl Zeiss NTS Supra 40 microscope.

### Supplementary Figures

**Supplementary Fig. 1.**
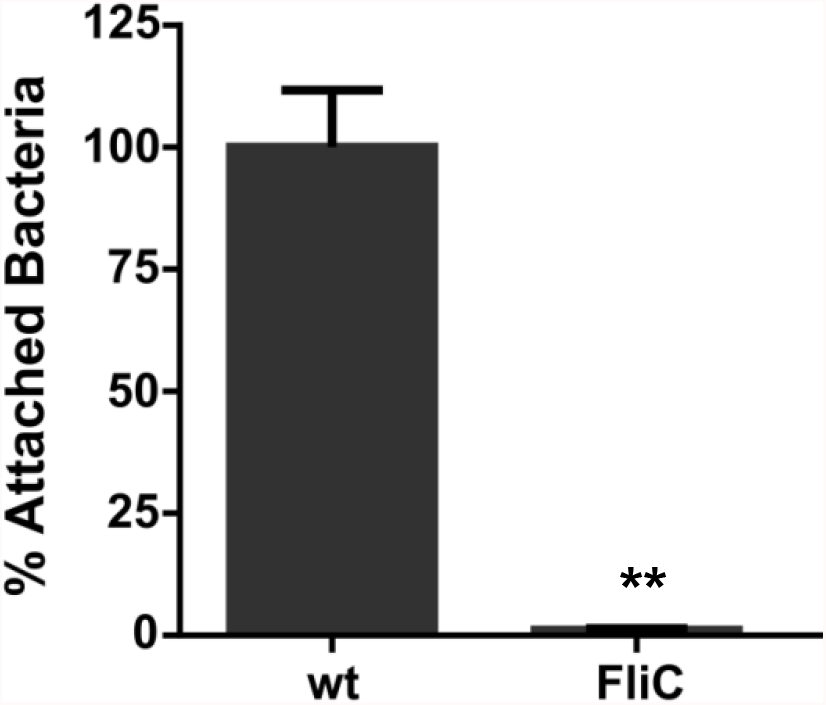
The flagellum is necessary for aggregate formation. To measure aggregate formation transwell grown MDCK monolayers were infected with wt P. aeruginosa-GFP and with ΔFliC-GFP, stained with Annexin V-Alexa 647 and fixed. The number of bacteria per aggregate was established using the ImageJ software as described by Lepanto et al. (*Mol. and Cell. Probes* 28, 1-5, 2014). Data are mean ± SEM. n = 3. ** p<0,01, Student’s *t*-test

**Supplementary Fig. 2.**
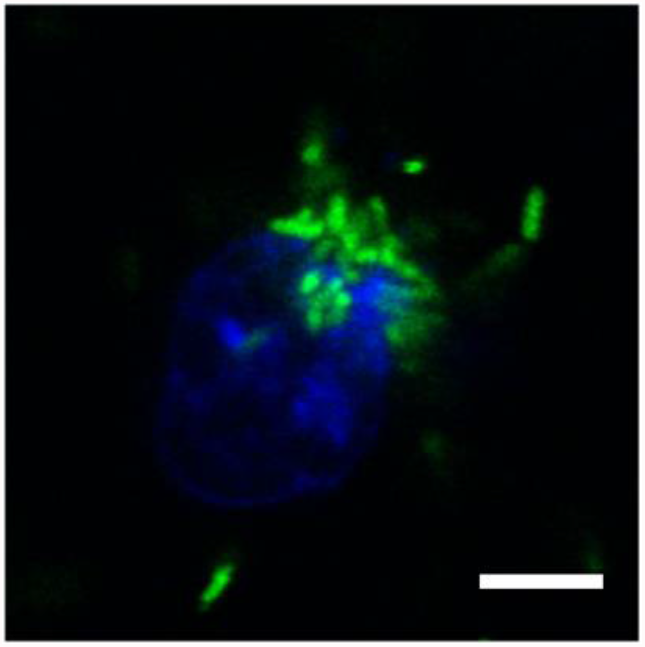
Bacteria adhere to zones of higher membrane surface availability. Time-lapse confocal image showing an extruded apoptotic cell with polarized bacterial adhesion. Prior to infection with *P. aeruginosa*-GFP (green), the monolayers were stained for 10 minutes with a general membrane marker (blue), (CellMask, Invitrogen). Bacterial binding occurred in zones where membrane labeling was more intense. Scale bar, 5 μm.

**Supplementary Fig. 3.**
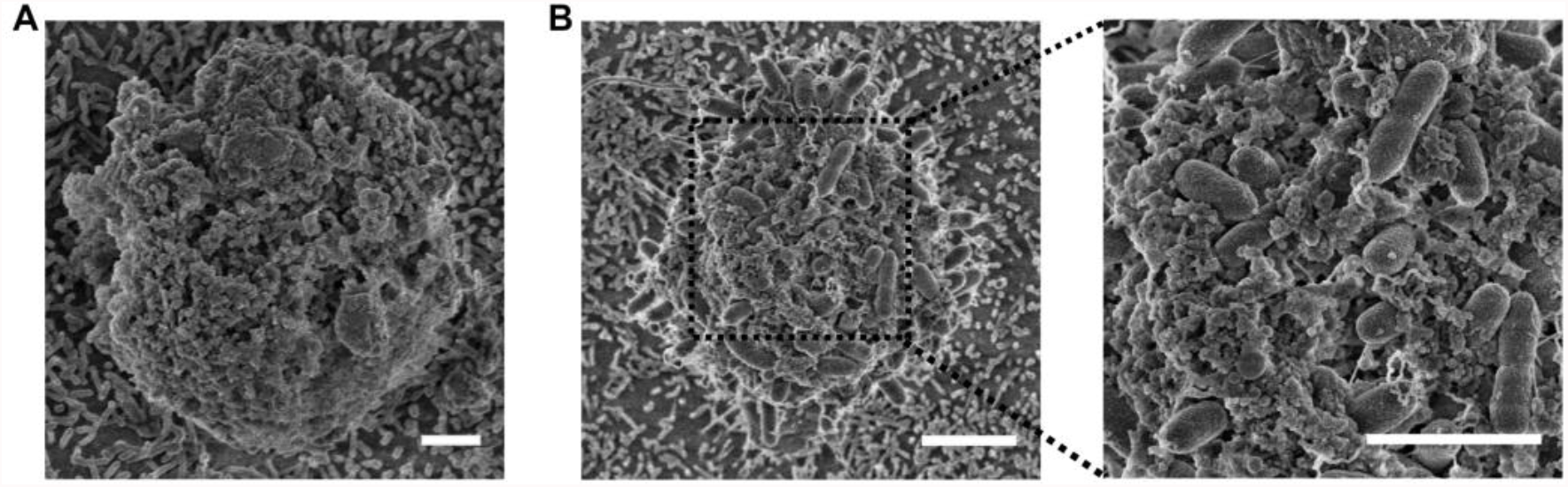
Bacteria adhere to apoptotic cells of vesiculated membrane morphology. Scanning electron microscopy images of uninfected and infected transwell grown MDCK monolayers. (A) The vesiculated membrane morphology is present in uninfected samples. (B) Monolayers were incubated for 1 h with *P. aeruginosa*. Extruded apoptotic cells homogeneously covered with bacteria are vesiculated all over their surface (left panel). Zoom in: Some bacteria are inserted between surface protuberances (right panel). Scale bars, 2 μm.

**Supplementary Fig. 4.**
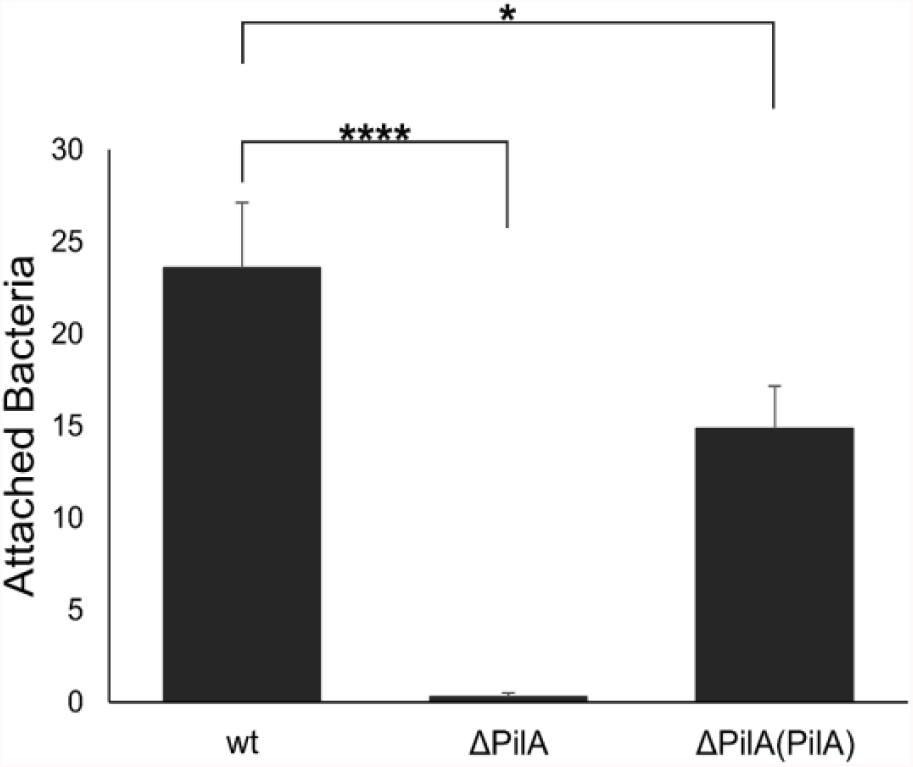
A non-piliated mutant is unable to aggregate on apoptotic cells. Transwell grown MDCK monolayers were infected for 1h with wt *P. aeruginosa*, the non-piliated mutant ΔPilA (the gen pila encodes for type four pili major subunit) and with the complemented mutant. To visualize bacteria samples were stained with the anti-pseudomonas antibody. The number of bacteria per aggregate was established using the ImageJ software as described by Lepanto et al. (*Mol. and Cell. Probes* 28, 1-5, 2014). Data are mean ± SEM, n = 3. *p< 0.05, ***p < 0.005, ****p < .0001, one-way ANOVA.

**Supplementary Fig. 5.**
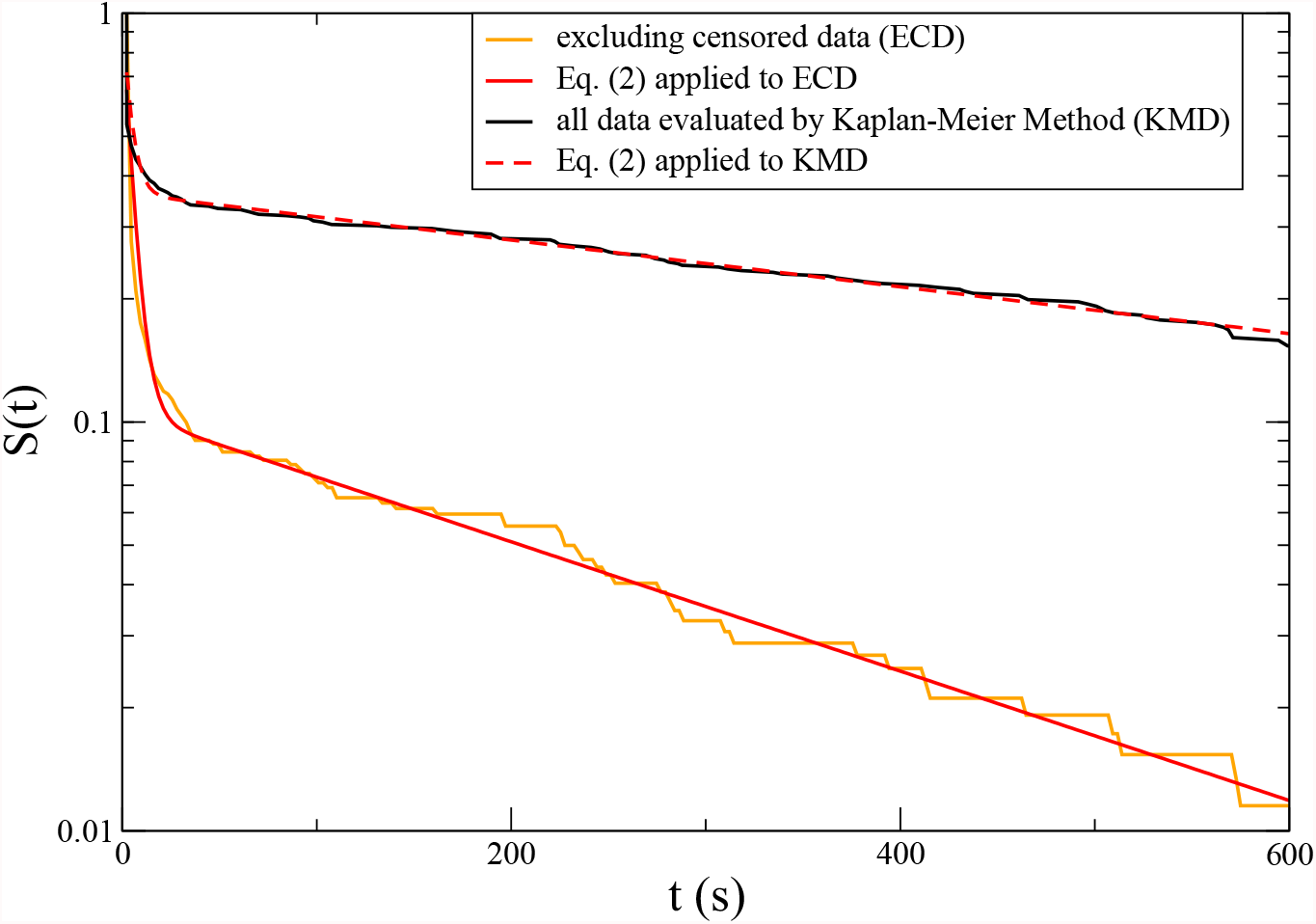
Cumulative probability distribution of WT residence times. Dwelling times are classified in two categories: those where we have observed the beginning and the end of the event (uncensored data), and those where we have observed the beginning, but not the end (censored data).

### Supplementary Movies

Supplemenatary Movie 1. Equatorial plane of an extruded apoptotic cell with heterogeneous AnnexinV staining (blue) infected with wt *P. aeruginosa* (green).

Supplementary Movie 2. Equatorial plane of two apoptotic cells (blue) infected with wt *P. aeruginosa* (green).

Supplementary Movie 3. Equatorial plane of apoptotic cell (blue) initially infected with wt P. aeruginosa-GFP (green) and 30 minutes later with wt P. aeruginosa-mCherry (red).

Supplementary Movie 4. Equatorial plane of apoptotic cell (blue) co-infected with wt *P. aeruginosa* (green) and the ΔPilA mutant (red).

Supplementary Movie 5. Equatorial plane of apoptotic cell (blue) co-infected with wt *P. aeruginosa* (red) and the ΔPilT mutant (green).

## References

1. Otte, S., Ipiña, E. P., Pontier-Bres, R., Czerucka, D. & Peruani, F. Statistics of pathogenic bacteria in the search of host cells. Nature communications 12, 1–9 (2021).

2. Bjarnsholt, T. et al. Pseudomonas aeruginosa biofilms in the respiratory tract of cystic fibrosis patients. Pediatric pulmonology 44, 547–558 (2009).

3. Burmølle, M. et al. Biofilms in chronic infections–a matter of opportunity– monospecies biofilms in multispecies infections. FEMS Immunology & Medical Microbiology 59, 324–336 (2010).

4. Bjarnsholt, T. et al. The in vivo biofilm. Trends in microbiology 21, 466–474 (2013).

5. Sønderholm, M. et al. Pseudomonas aeruginosa aggregate formation in an alginate bead model system exhibits in vivo-like characteristics. Applied and environmental microbiology 83 (2017).

6. Jennings, L. K. et al. Pseudomonas aeruginosa aggregates in cystic fibrosis sputum produce exopolysaccharides that likely impede current therapies. Cell reports 34, 108782 (2021).

7. Karygianni, L., Ren, Z., Koo, H. & Thurnheer, T. Biofilm matrixome: extracellular components in structured microbial communities. Trends in Microbiology 28, 668–681 (2020).

8. Wong, G. C. et al. Roadmap on emerging concepts in the physical biology of bacterial biofilms: from surface sensing to community formation. Physical biology 18, 051501 (2021).

9. Armbruster, C. R. & Parsek, M. R. New insight into the early stages of biofilm formation. Proceedings of the National Academy of Sciences 115, 4317–4319 (2018).

10. Berne, C., Ellison, C. K., Ducret, A. & Brun, Y. V. Bacterial adhesion at the single-cell level. Nature Reviews Microbiology 16, 616–627 (2018).

11. Lee, C. K. et al. Multigenerational memory and adaptive adhesion in early bacterial biofilm communities. Proceedings of the National Academy of Sciences 115, 4471–4476 (2018).

12. Lee, C. K. et al. Social cooperativity of bacteria during reversible surface attachment in young biofilms: A quantitative comparison of pseudomonas aeruginosa pa14 and pao1. Mbio 11 (2020).

13. Wu, S., Zhang, B., Liu, Y., Suo, X. & Li, H. Influence of surface topography on bacterial adhesion: A review. Biointerphases 13, 060801 (2018).

14. Song, F., Koo, H. & Ren, D. Effects of material properties on bacterial adhesion and biofilm formation. Journal of dental research 94, 1027–1034 (2015).

15. Sauer, K., Camper, A. K., Ehrlich, G. D., Costerton, J. W. & Davies, D. G. Pseudomonas aeruginosa displays multiple phenotypes during development as a biofilm. Journal of bacteriology 184, 1140–1154 (2002).

16. Davey, M. E., Caiazza, N. C. & O’Toole, G. A. Rhamnolipid surfactant production affects biofilm architecture in pseudomonas aeruginosa pao1. Journal of bacteriology 185, 1027–1036 (2003).

17. Lepanto, P. et al. Pseudomonas aeruginosa interacts with epithelial cells rapidly forming aggregates that are internalized by a lyn-dependent mechanism. Cellular microbiology 13, 1212–1222 (2011).

18. Capasso, D. et al. Elimination of pseudomonas aeruginosa through efferocytosis upon binding to apoptotic cells. PLoS pathogens 12, e1006068 (2016).

19. Denise, R., Abby, S. S. & Rocha, E. P. Diversification of the type iv filament superfamily into machines for adhesion, protein secretion, dna uptake, and motility. PLoS biology 17, e3000390 (2019).

20. Chang, Y.-W. et al. Architecture of the type iva pilus machine. Science 351 (2016).

21. Caiazza, N. C. & O’Toole, G. A. Sadb is required for the transition from reversible to irreversible attachment during biofilm formation by pseudomonas aeruginosa pa14 (2004).

22. Güvener, Z. T., Tifrea, D. F. & Harwood, C. S. Two different pseudomonas aeruginosa chemosensory signal transduction complexes localize to cell poles and form and remould in stationary phase. Molecular microbiology 61, 106–118 (2006).

23. Murray, T. S. & Kazmierczak, B. I. Flhf is required for swimming and swarming in pseudomonas aeruginosa. Journal of bacteriology 188, 6995–7004 (2006).

24. Cowles, K. N. et al. The putative poc complex controls two distinct p seudomonas aeruginosa polar motility mechanisms. Molecular microbiology 90, 923–938 (2013).

25. Thomas, W. E., Vogel, V. & Sokurenko, E. Biophysics of catch bonds. Annu. Rev. Biophys. 37, 399–416 (2008).

26. Koch, M. D., Fei, C., Wingreen, N. S., Shaevitz, J. W. & Gitai, Z. Competitive binding of independent extension and retraction motors explains the quantitative dynamics of type iv pili. Proceedings of the National Academy of Sciences 118 (2021).

27. Schniederberend, M. et al. Modulation of flagellar rotation in surface-attached bacteria: a pathway for rapid surface-sensing after flagellar attachment. PLoS pathogens 15, e1008149 (2019).

28. Ipiña, E. P., Otte, S., Pontier-Bres, R., Czerucka, D. & Peruani, F. Bacteria display optimal transport near surfaces. Nature Physics 15, 610–615 (2019).

29. Pierrat, X., Wong, J. P., Al-Mayyah, Z. & Persat, A. The mammalian membrane microenvironment regulates the sequential attachment of bacteria to host cells. Mbio 12, e01392–21 (2021).

30. Hehemann, J.-H. et al. Adaptive radiation by waves of gene transfer leads to fine-scale resource partitioning in marine microbes. Nature communications 7, 1–10 (2016).

31. Wargacki, A. J. et al. An engineered microbial platform for direct biofuel production from brown macroalgae. Science 335, 308–313 (2012).

32. Ratzke, C. & Gore, J. Self-organized patchiness facilitates survival in a cooperatively growing bacillus subtilis population. Nature microbiology 1, 1–5 (2016).

